## Supplementary figures and images for "E-cadherin endocytosis promotes non-canonical EGFR:STAT signalling to induce cell death and inhibit heterochromatinisation"

### Supplemental Figures

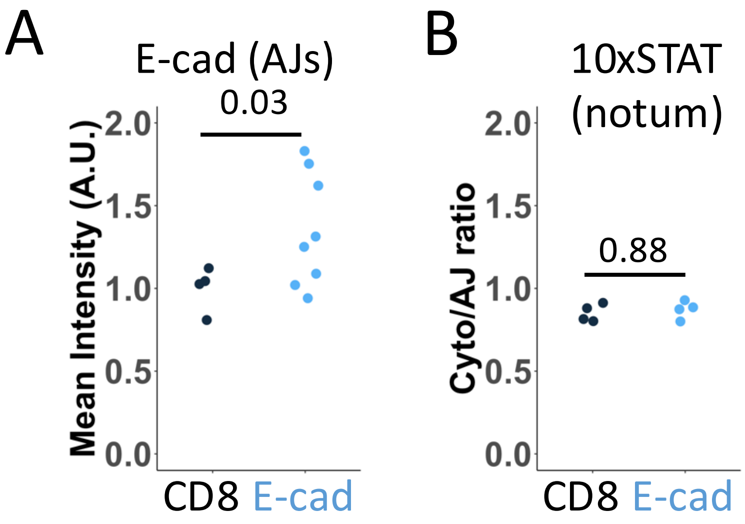

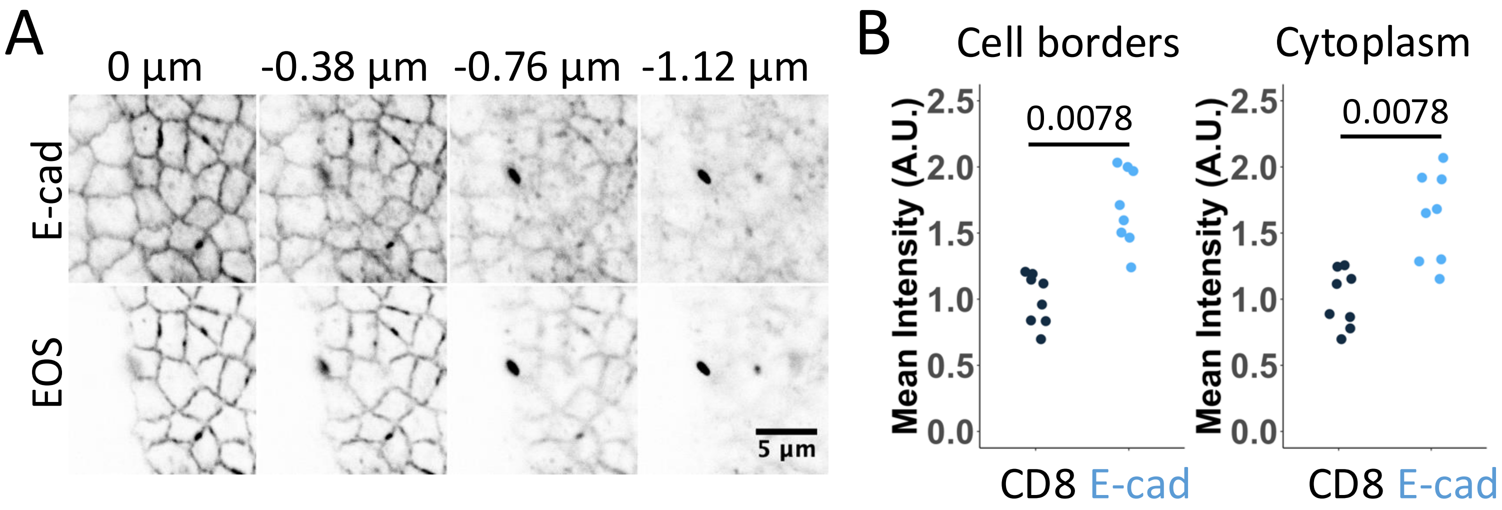

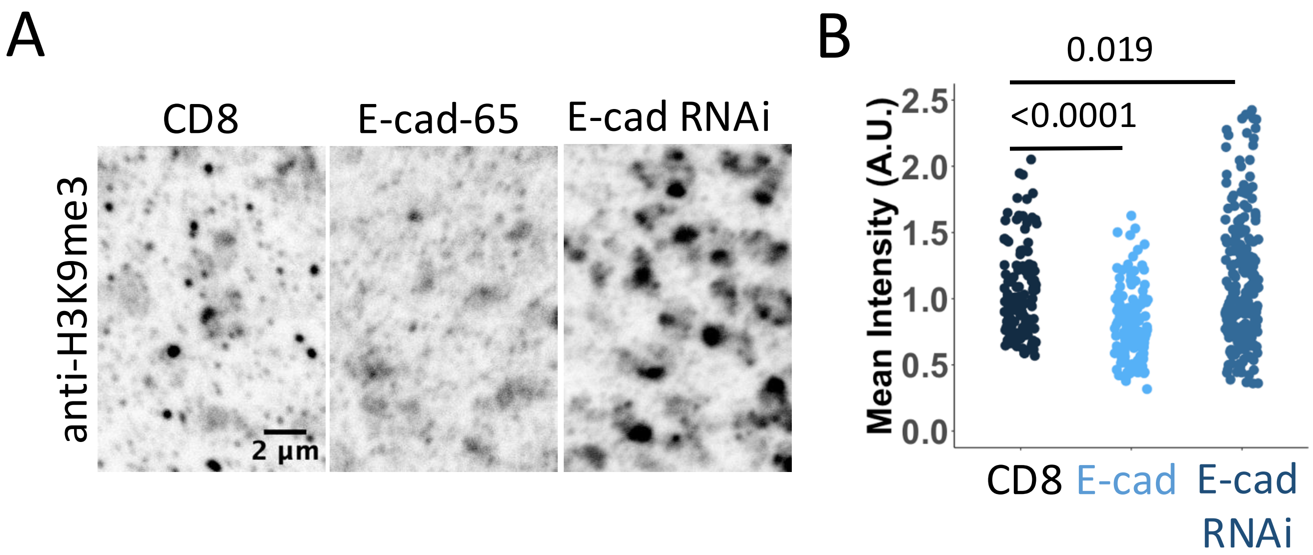

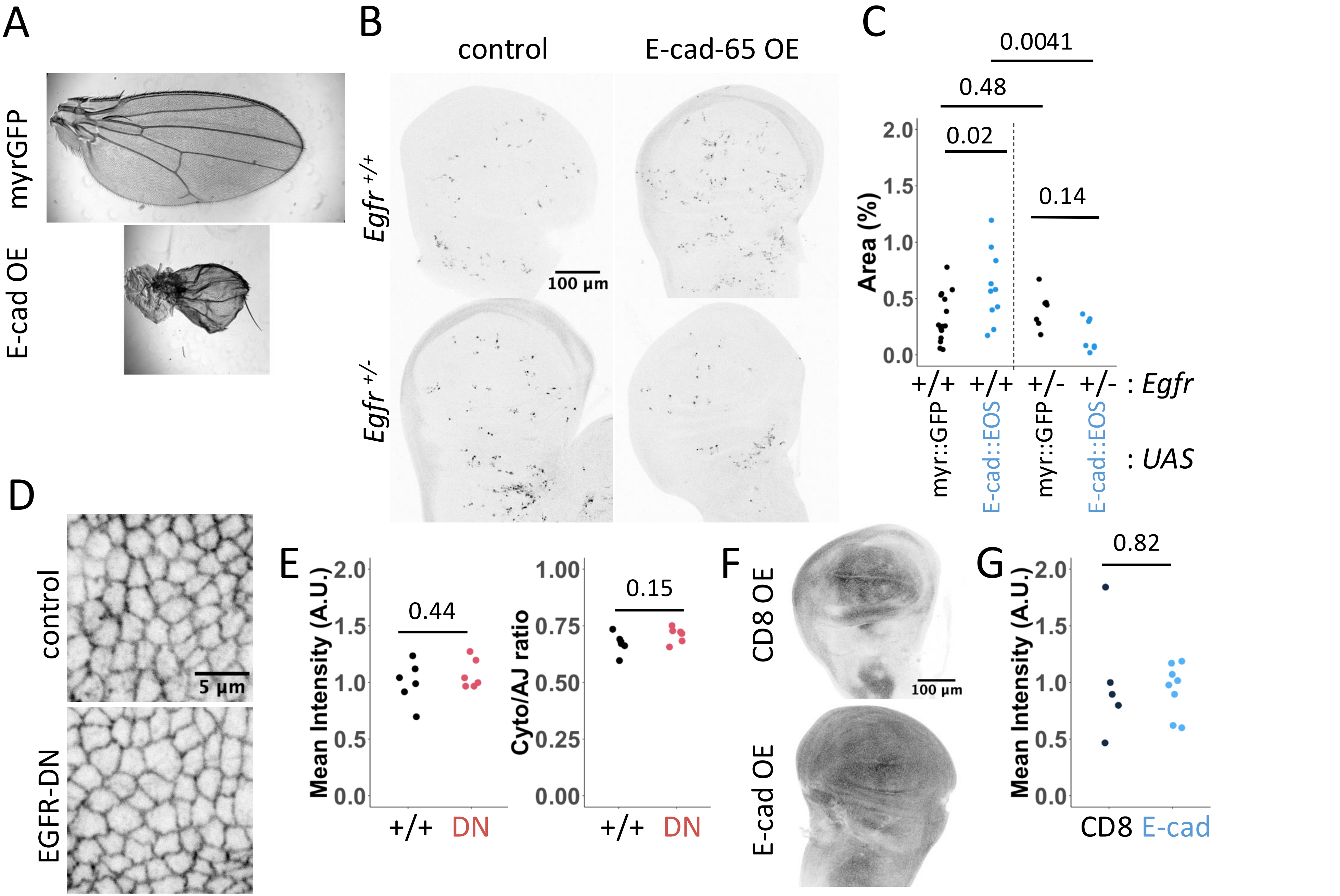

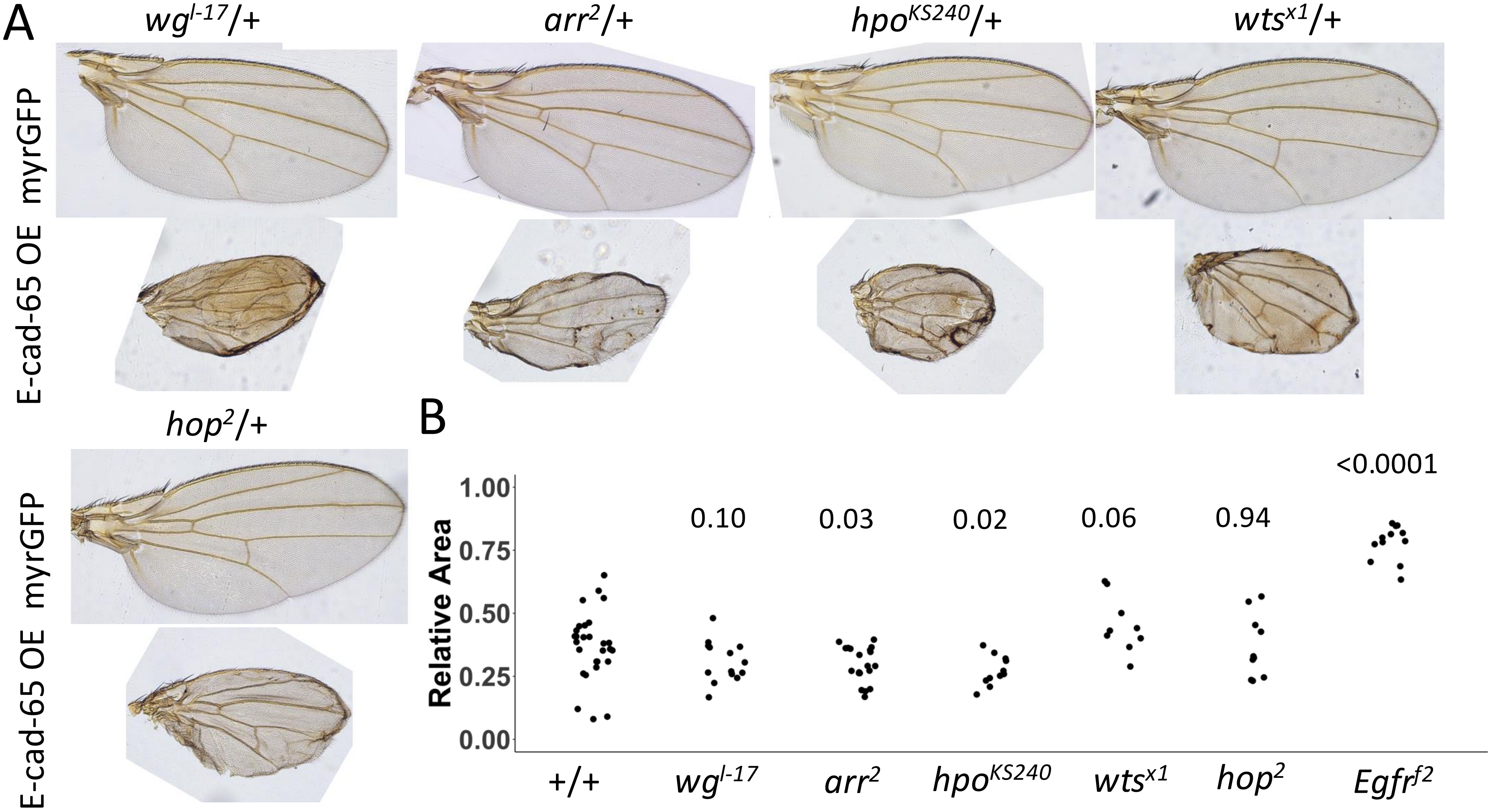
